# Lipopolysaccharide-induced systemic inflammation alters courtship ultrasonic vocalizations in male mice

**DOI:** 10.64898/2026.04.13.718006

**Authors:** Mizuki Yamamoto, Haruki Kasahara, Kazuko Hayashi, Yuta Tamai, Koji Toda

**Affiliations:** Department of Psychology, Keio University, Tokyo, JAPAN; Japan Society for the Promotion of Science, Tokyo, JAPAN; Department of Otolaryngology-Head and Neck Surgery, Hearing Research Centre, University of Tübingen, Tübingen, GERMANY

**Keywords:** ultrasonic vocalization, communication, social interaction, sickness behavior, lipopolysaccharide, inflammation, mice

## Abstract

**Background:** Sickness behavior comprises a coordinated constellation of motivational, cognitive, and social alterations that emerge during systemic inflammation. Although reductions in locomotion, feeding, and social engagement have been extensively characterized, how inflammation affects ultrasonic vocal communication—an ethologically relevant index of social motivation in rodents—remains insufficiently understood. Here, we investigated how systemic immune activation alters male–female social communication in mice by jointly assessing ultrasonic vocalizations (USVs) and approach behavior.

**Methods:** Sexually experienced male mice received an intraperitoneal injection of lipopolysaccharide (LPS), and their interactions with a novel estrous female were evaluated 24 h later by quantifying USVs and approach behavior.

**Results:** LPS administration robustly suppressed both the total number of USVs and the duration of male approach behavior, indicating a pronounced reduction in social motivation. Beyond this quantitative suppression, LPS also induced qualitative changes in vocal output, including shifts in the proportional use of specific USV subtypes and alterations in acoustic features such as sound pressure.

**Conclusions:** These findings demonstrate that USVs capture multiple dimensions of inflammation-induced disruption of social communication, reflecting not only diminished motivation to engage socially but also changes in the structure of communicative signals themselves. By revealing that systemic immune activation reshapes both social approach behavior and vocal communication patterns, this study establishes USV analysis as a sensitive and translationally relevant behavioral readout for probing neuroimmune mechanisms underlying the social and communicative disturbances characteristic of sickness behavior. More broadly, our results highlight the utility of vocal communication analyses for elucidating how inflammatory processes perturb social circuits and communicative function in health and disease.

## 1. Introduction

Sickness behavior refers to a coordinated constellation of behavioral, motivational, and social alterations that emerge during systemic infection or inflammation. In animals, this state is characterized by changes in sleep and circadian activity, reduced food and water intake, hypoactivity, and marked disruptions in social engagement (Dantzer et al., 2023). These responses, often accompanied by fever and lethargy, are widely regarded as an evolutionarily conserved adaptive strategy that reallocates energy toward immune defense while limiting pathogen transmission (Kluger et al., 1975).

Experimentally, sickness behavior is most commonly modeled through peripheral administration of lipopolysaccharide (LPS), a bacterial endotoxin that activates Toll-like receptor 4 (TLR4)–dependent innate immune signaling. LPS administration triggers robust cytokine release and engages multiple pathways through which peripheral immune activation influences central nervous system function, including humoral routes, vagal afferents, and cytokine transport across the blood–brain barrier (Lasselin, 2021; Lu et al., 2008; Park & Lee, 2013). Consistent with this framework, LPS reliably induces reductions in locomotion, food intake, body weight, hedonic processing, and a broad range of social behaviors in rodents (Clark et al., 2015; Lasselin et al., 2020; Osterhout et al., 2022; Oliveira et al., 2024). In particular, prior studies have documented decreased social exploration (Bluthé et al., 1994; Yee et al., 2010), impaired maternal care (Aubert et al., 1997; Bernardi et al., 2009), and attenuated sexual behavior following inflammatory challenge (Yirmiya, 1996).

Despite this extensive behavioral literature, most studies have relied on coarse measures such as time spent in proximity, mounting frequency, or overall activity levels, which provide limited insight into the quality and structure of social communication itself. One especially underexplored domain is ultrasonic vocal communication. Ultrasonic vocalizations (USVs) constitute a primary communicative signal through which mice actively transmit motivational and affective states to conspecifics across diverse social contexts, including courtship, maternal bonding, territorial interactions, and affiliative contact (Premoli et al., 2022; Ehret, 2005; Gourbal et al., 2004). Importantly, USVs provide a non-invasive and temporally precise behavioral readout that can capture internal state changes not readily inferred from overt motor behavior alone.

Accumulating evidence suggests that USVs are highly sensitive to changes in emotional and physiological states across a wide range of disease models (Premoli et al., 2023). Altered USV patterns have been reported in models of autism spectrum disorders, neurodevelopmental disruption, neurodegeneration, chronic stress, and pain, often preceding or occurring independently of gross motor impairments. For example, chronic cancer pain and neuropathic pain markedly increase USV emission within specific frequency bands, particularly around 50 kHz, reflecting ongoing spontaneous pain states rather than stimulus-evoked motor responses (Kurejova et al., 2010). These findings highlight USVs as a powerful and ethologically relevant metric for assessing subtle changes in internal motivational and affective states.

A particularly informative context in which to examine inflammation-induced modulation of USVs is male–female interaction. During heterosexual encounters, male mice produce the vast majority of USVs, and these vocalizations exert a direct influence on female approach behavior and mate choice (Willadsen et al., 2014). Females preferentially approach males that emit more frequent, sustained, and structurally complex USVs, indicating that vocal output serves as a reliable proxy for sexual motivation and communicative competence (Tschida et al., 2019). Importantly, male courtship USVs are tightly regulated by neural circuits that integrate motivational, affective, and motor components of social behavior, making them particularly vulnerable to disruption by inflammatory signaling.

Although systemic inflammation is known to suppress sexual behavior and social motivation, it remains unclear whether such effects extend to the structure and quality of social communication signals themselves. Reductions in USV emission could arise secondarily from decreased locomotion or physical proximity, or alternatively reflect a more specific impairment in the neural mechanisms governing socio-communicative drive and vocal production. Distinguishing between these possibilities is essential for understanding whether sickness behavior merely dampens behavioral output globally or selectively alters communicative processes.

Based on this framework, the primary aim of the present study was to determine whether systemic inflammation suppresses male ultrasonic vocal communication during male–female interactions. While it is well established that inflammatory challenges induce sickness behavior and reduce sexual motivation (Dantzer et al., 2008), it remains unclear whether inflammation specifically disrupts vocal communication itself, beyond generalized reductions in social or locomotor activity. To address this question, we first assessed male approach behavior toward a novel estrous female to confirm the expected reduction in sexual motivation following LPS administration and to establish the behavioral context in which vocalizations occur. We then examined whether LPS-induced inflammation decreases the total number of male USVs emitted during courtship interactions while controlling for spatial proximity and locomotor activity. Finally, using machine-learning–based acoustic analysis (Fonseca et al., 2021), we tested whether inflammation alters the fine-grained spectral organization of USVs beyond changes in vocal output alone, enabling an unbiased assessment of inflammation-induced disruptions in social communication. Importantly, our goal was not simply to confirm a reduction in sexual motivation, but to test whether systemic inflammation selectively impairs vocal communication as a component of social signaling.

## 2. Methods

### 2.1. Subjects

All data were collected from 2.3- to 4.5-month-old adult male (average: 2.7, SD: 0.64) and 2.3- to 4.4-month-old adult female wild-type C57BL/6J mice (average: 3.5, SD: 0.54). The animals were maintained on a 12:12 h light/dark cycle, and all experiments were conducted during the dark phase. Mice had ad libitum access to food and water in their home cages. All animal procedures were conducted in accordance with the National Institutes of Health Guide for the Care and Use of Laboratory Animals and were approved by the Animal Research Committee of Keio University.

### 2.2. Drug

Lipopolysaccharide (LPS; CAS L2630, Sigma-Aldrich, Gillingham, UK) was dissolved in saline, divided into small aliquots, and stored at −80 °C. LPS was administered intraperitoneally at a dose of 1.0 mg/kg (0.01 mL/g) in all experiments. Regarding the timing of administration, all mice received saline injection 24 h before Experiment 1. For Experiment 2, mice received either saline or LPS (1.0 mg/kg) intraperitoneally 24 h before behavioral testing. The LPS dose used in this study (1.0 mg/kg) was selected based on doses frequently used in previous studies (0.5, 0.83, and 1.0 mg/kg), which together accounted for 73.19% of studies in which LPS was administered intraperitoneally to mice (Yin et al., 2023). These doses have been reported to increase immobility time in the forced swim test and decrease in locomotor activity in C57BL/6 mice (Kong et al., 2021; Morimoto et al., 2023; Yin et al., 2023). To ensure a consistent physiological response to LPS, mice that exhibited less than a 5% reduction in body weight following LPS injection were excluded from the analysis. This criterion was applied to exclude animals that were considered non-responders or showed an insufficient systemic response to LPS.

### 2.3. Procedure

#### 2.3.1. Habituation

We used a male-female social interaction task to examine the effects of intraperitoneal LPS injection on USVs and approach behavior of male mice toward female mice. Adult male mice were paired with an adult female mouse in a standard cage for 2–5 days prior to testing to ensure sexual experience. The duration of cohabitation varied within this range due to logistical constraints but did not differ systematically between experimental groups. This period was sufficient to provide sexual experience and was not intended to manipulate the degree of sexual experience across individuals. Males were then habituated to the experimental cage (width 28.0 cm × depth 9.5 cm × height 30.0 cm) for 1–2 days.

#### 2.3.2. Experiment 1: Recording the number of USVs emitted to an empty basket and a basket containing a novel female mouse

Following habituation, we first confirmed whether male mice emitted USVs as a courtship behavior. We conducted two conditions: (1) an empty basket and (2) a basket containing a novel female mouse. USVs and approach behavior were recorded for 10 min under each condition (Figure 1A). The two conditions were presented in a counterbalanced order for potential order effects, such as habituation and novelty. Thus, some subjects were exposed first to the female stimulus followed by the empty basket, whereas others experienced the opposite order. Both recordings were conducted on the same day. In this experiment, males received an intraperitoneal injection of saline 5 min before testing. Subject males were then temporarily removed from the experimental cage to allow placement of the basket. To prevent climbing, a 500-mL empty glass bottle was placed on top of the basket. Immediately after returning the subjects to the cage, audio and video recording was initiated. Audio and video signals were synchronized using a custom-made Arduino sketch that generated a brief beep and LED flash simultaneously at the onset and offset of recording. Audio data were analyzed immediately after the experiment, and only males that produced USVs in the female-present condition were included in subsequent Experiment 2. Males producing USVs were operationally defined as those emitting more than 100 USVs during the 10-min courtship condition. This criterion was based on the distribution of USV counts observed in the dataset: males emitting fewer than 100 USVs showed little to no vocal output, whereas most males exceeding this threshold typically produced approximately 500 USVs in total. This cutoff was therefore applied to exclude males with negligible USV production from further analyses.

**Figure 1.**
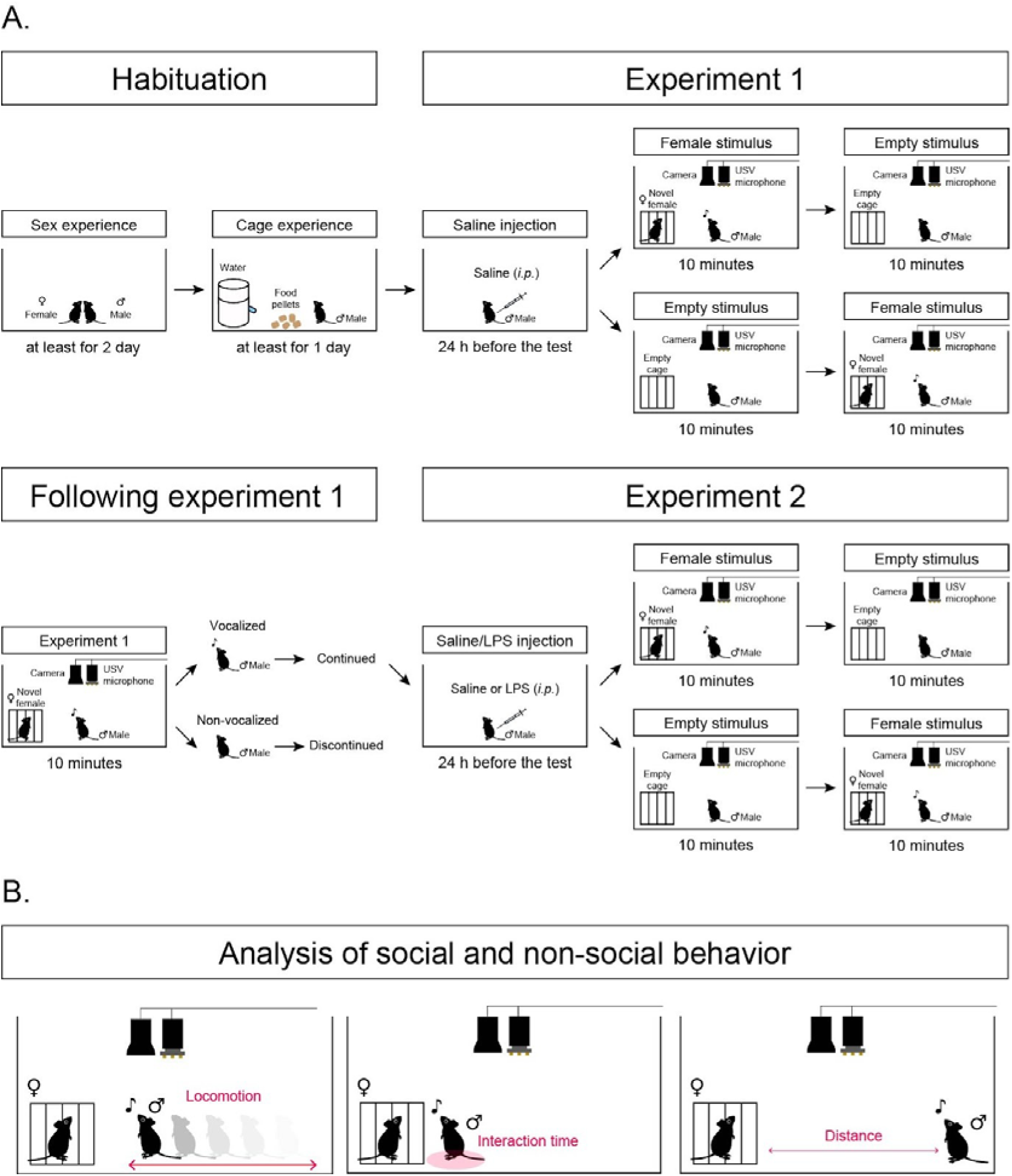
A. Schematic diagram of the social interaction test procedure. Habituation: Male mice were paired with a female mouse in their home cage for 2-5 days prior to testing to ensure sexual experience. Following this pairing period, each male mouse was habituated to the experimental cage for 1-2 days. Experiment 1: After habituation, male mice received a saline injection 24 h before testing. During the test session, each mouse was exposed to either a novel estrous female mouse enclosed in a wire basket (female stimulus condition) or an empty basket (empty-basket condition), presented in a counterbalanced order. In the female stimulus condition, only males that emitted USVs were included in subsequent experiments; mice that did not vocalize were excluded from further testing. Experiment 2: Male mice received an intraperitoneal injection of either saline or LPS 24 h before testing. Twenty-four hours after injection, each mouse was exposed to a novel female mouse or an empty basket in a counterbalanced order, following the same testing procedure as in Experiment 1. B. Schematic diagram of the analysis of the social interaction test.

#### 2.3.3. Experiment 2: Recording the number of USVs emitted to a basket containing a novel female mouse after saline or LPS treatment

On the day after the experiment 1, vocalizing males received an intraperitoneal injection of LPS (1.0 mg/kg) or saline 24 h before the experiment 2. During the Experiment 2 test following the injection, a novel female mouse was introduced into the basket, and USVs and approach behavior were recorded for 10 min using the same experimental setup as in the experiment 1 (Figure 1A).

A condenser microphone (CM16/CMPA, Avisoft Bioacoustics) was positioned 31 cm above the floor of the experimental box to record USVs. The recorded sound data was stored on a computer through an audio interface (UltraSoundGate 116Hb, Avisoft Bioacoustics) using a sampling rate of 96 kHz at 16 bits/sample. A camera (C980GR, Logicool) was placed next to the microphone and used to track the location of the subject male mouse. All experiments were conducted inside a sound-attenuating chamber (ENV-018V, Med Associates Inc.) to minimize external noise. After each session, inside of the experimental box, basket, and water bottle were wiped with a paper towel soaked with 70% ethanol and allowed to air-dry for 30 min before reuse.

### 2.4. Analysis

All data analyses were performed using RStudio (version 2022.02.0, RStudio PBC, MA, USA), MATLAB (R2020a, The MathWorks, Inc., Natick, MA, USA), Python (version 3.8.10), and GraphPad Prism (version 10.3.1, GraphPad, CA, USA). Locomotor activity and social interaction of mice was tracked using the open-source visual programming framework Bonsai (Lopes et al., 2015) for computer vision analysis (Figure 1B). Video recordings were converted to grayscale, smoothed, and inverted to enhance contrast, and mice were detected by applying a contrast threshold. Locomotor activity and locations within the experimental box was quantified by calculating changes in the animals’ central coordinates. We defined mice as spending time with the social interaction if they were located within 3.5 cm of the edge of the box that contained the stimulus mice.

To investigate the characteristics of USVs, VocalMat (Fonseca et al., 2021), a machine learning-based platform that enables segmentation of the USVs from the background noise and classifies USVs into distinct call categories, was used. After the segmentation of USVs via VocalMat, the spectrotemporal features of USVs were quantified using the following parameters with a custom-made MATLAB script: call duration, dominant frequency, frequency range (maximum-minimum), and Wiener entropy. For each USV syllable, spectrograms were computed using Fast Fourier Transform (1024 points with a Hanning window). The relative sound pressure level of the F0 component was calculated in decibels by normalizing each vocal call to the call with the maximum signal amplitude; the loudest call was assigned 0 dB. Acoustical feature analysis was conducted based on the framework described in previous studies (Kobayasi and Riquimaroux, 2012; Furuyama et al., 2022), with modifications to investigate more precise acoustic features.

Dimensionality reduction of these acoustic features was performed using Uniform Manifold Approximation and Projection (UMAP) to assess the internal relationship among the different parameters of simple syllables. Prior to the UMAP analysis, all acoustic features were standardized to a zero mean and unit variance. The distribution of calls in the latent space was compared in each experimental condition (Saline vs LPS). The UMAP analysis was performed with a custom-made Python script.

## 3. Results

### 3.1. Experiment 1

To investigate the relationship between male–female social interactions and USVs, we examined male–female USVs as well as social and non-social behaviors using the social interaction test in Experiment 1. To verify that the observed USVs reflected courtship-related behavior, we included an empty-stimulus condition, in which a basket without a female mouse was presented. Male mice were exposed to two conditions: a female stimulus condition, in which an estrous female mouse was placed inside a basket, and an empty-basket condition, in which the basket was presented without a female (Figure 1A). USVs were analyzed using VocalMat (Fonseca et al., 2021), which successfully classified vocalizations into 11 categories—chevron, reverse chevron, down-frequency modulation, up-frequency modulation, flat, short, complex, step-up, step-down, two steps, and multiple steps—and discriminated them from noise (Figure 2). We first examined whether the subject mice emitted USVs in response to the presentation of a female mouse. This initial screening revealed that 26 out of 37 mice (70.3%) produced USVs in response to the female stimulus (vocalizing mice, Figure S1A). Vocalizing mice were older than non-vocalizing mice (Figure S1B; *t*(33.6) = 2.60, *p* = .0137, unpaired t-test). However, the number of USVs was not significantly correlated with age (Figure S1C; *r* = 0.3208, *p* = .0909, Pearson’s correlation). Body weight did not differ between vocalizing and non-vocalizing mice (Figure S1D; *t*(31.0) = 1.56, *p* = .128, unpaired t-test). In addition, the number of the USVs was not significantly correlated with body weight (Figure S1E; *r* = 0.2781, *p* = .0528, Pearson’s correlation). Locomotor activity showed a significant condition × group interaction (Figure S1F, G; condition x group interaction: *F*(1, 35) = 10.25, *p* = .0029, two-way ANOVA). Post hoc Fisher’s LSD tests revealed that locomotor activity was higher in the female condition than in the empty condition in vocalizing mice (*p* = .0445), whereas the opposite pattern was observed in non-vocalizing mice (*p* = .0188). In addition, in the empty condition, locomotor activity was higher in vocalizing mice than in non-vocalizing mice (*p* = .0014). Approach behavior toward the female stimulus did not differ between vocalizing and non-vocalizing subjects (Figure S1H–K; interaction time: all *p*s > .05; distance: all *p*s > .05, two-way ANOVA). We further examined whether stimulus presentation order affected USV emission. A repeated-measures two-way ANOVA revealed a significant main effect of stimulus (Figure S2; *F*(1, 42) = 44.59, *p* < .0001), but no effect of presentation order or interaction (stimulus presentation order: *F*(1, 42) = 0.7135, *p* = .4030; stimulus x stimulus presentation order interaction: *F*(1, 42) = 1.430, *p* = .2385). These results indicate that the observed differences in USV emission were not confounded by stimulus presentation order. The 15 mice that did not emit USVs (non-vocalizing mice) were excluded from subsequent analyses (Figure 3A). Male mice that emitted robust USVs in the presence of the female stimulus showed almost no USVs under the empty-basket condition (Figure 3B). Quantitative analysis confirmed that USV emission was significantly higher in the female stimulus condition than in the empty-basket condition (*t*(25) = 9.72, *p* < 0.0001, paired t-test). Analysis of USV emission across successive time bins demonstrated that mice consistently produced USVs throughout the session in the female stimulus condition, indicating that the presence of the female stimulus elicited persistent USV production over time (Figure 3C). We further examined the relationship between USV emission and the distance from the female stimulus. This analysis demonstrated that USV emission was significantly higher when mice were located closer to the female stimulus, indicating that vocal production was spatially coupled with social approach behavior (Figure 3E). Furthermore, analysis of USV categories as a function of both distance from the female stimulus and elapsed time since stimulus presentation showed no preferential emission of specific categories, indicating that neither spatial proximity to the female nor temporal factors biased the types of vocalizations produced (Figure 3D, F). In the social behavior, paired t-tests revealed that there were no significant differences in locomotor activity (Figure 3H, I; *t*(25) = 2.029, *p* = .0532), interaction time (Figure 3J, K; *t*(25) = 1.667, *p* = .1080) or mean distance from the stimulus (Figure 3L, M; *t*(25) = 2.029, *p* = .0532) between the female stimulus condition and empty-basket conditions.

**Figure 2.**
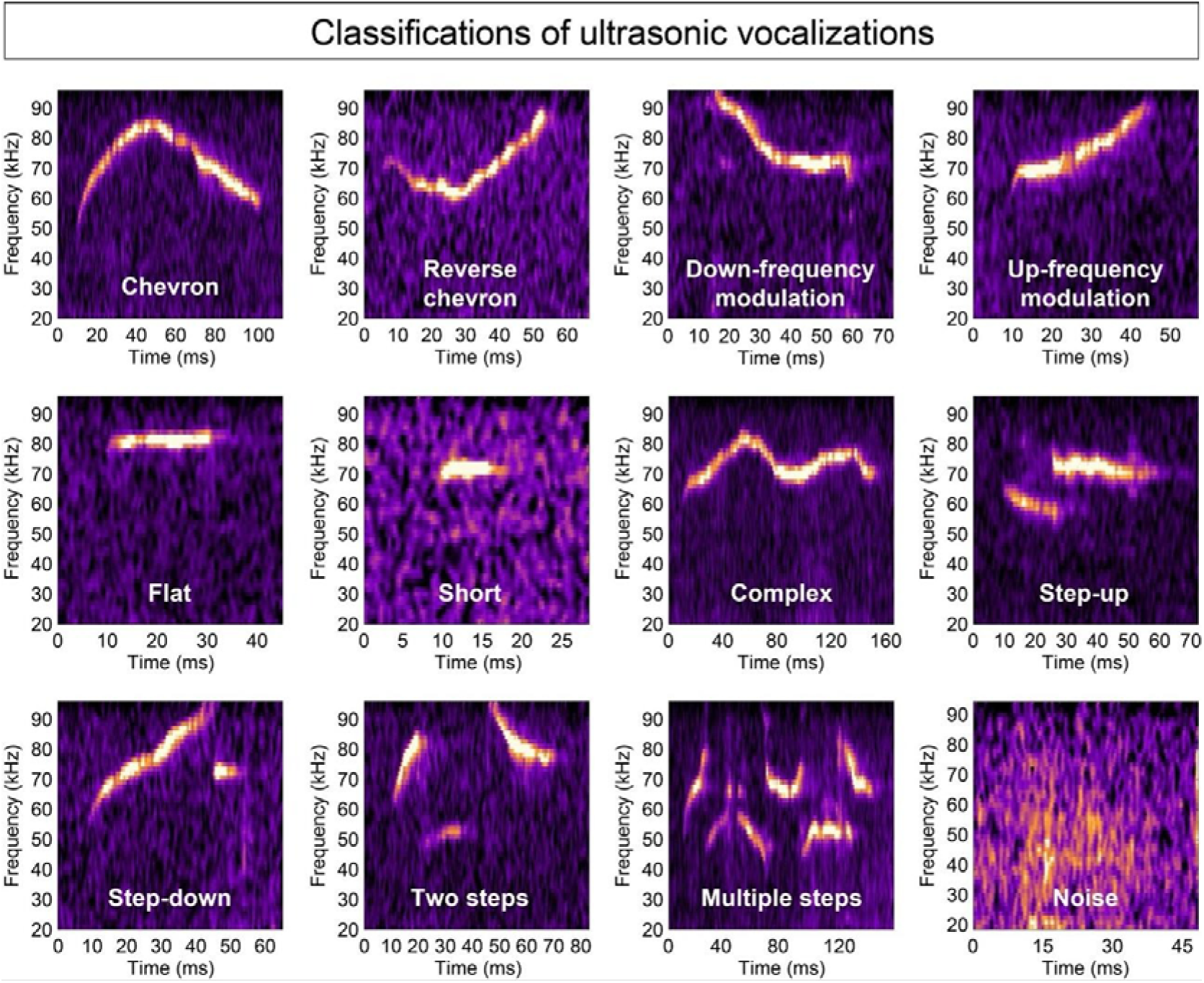
Example of the 11 categories of USVs and noise. USVs were analyzed using VocalMat, which successfully classified vocalizations into 11 categories—chevron, reverse chevron, down-frequency modulation, up-frequency modulation, flat, short, complex, step-up, step-down, two steps, and multiple steps—and discriminated them from noise.

**Figure 3.**
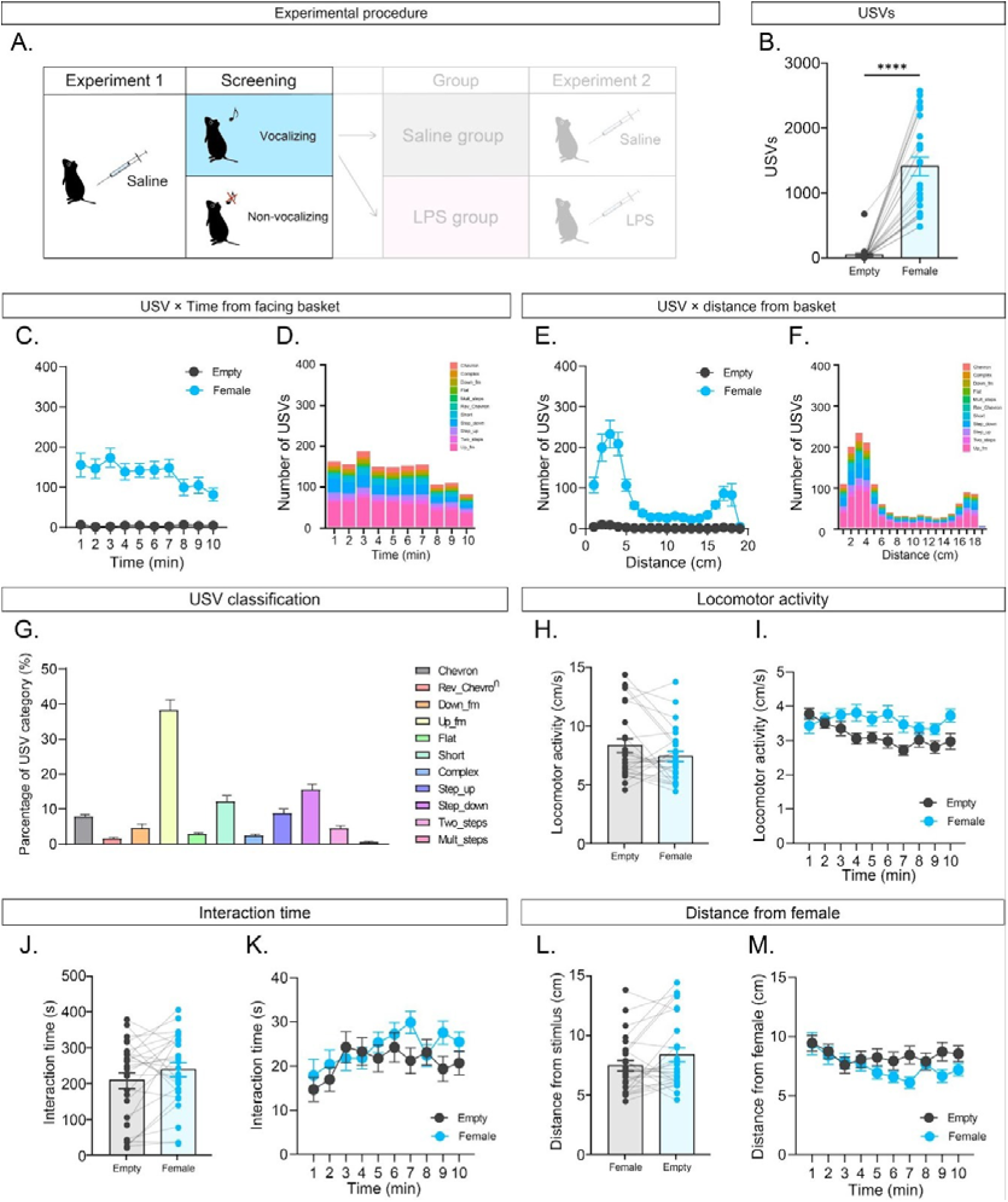
A. All mice were injected with saline in Experiment 1. Only vocalizing subjects were selected and assigned to two groups for subsequent saline or LPS administration in Experiment 2. B. Comparison of the number of USVs in the female stimulus condition and the empty condition. C. Time course of the number of USVs. D. Time course of the distribution of USV categories under the female condition. Comparison of number of USVs between the female stimulus and empty condition in vocalizing subjects. E.The number of USVs at different distances from the basket. F Time course of the distribution of USV categories at different distances from the basket under the female condition. G Distribution of USV categories under the female condition. H. Comparison of locomotor activity between the female and the empty condition. I. Time course of the locomotor activity. J. Comparison of social interaction time between the female and the empty condition. K. Time course of the social interaction time. L. Comparison of the distance from the basket between the female and the empty condition. M. Time course of the distance from the basket. Error bars indicate the standard error of the mean, \**p* < 0.05, ** *p* < 0.01, **** *p* < 0.0001, N = 13.

**Figure S1.**
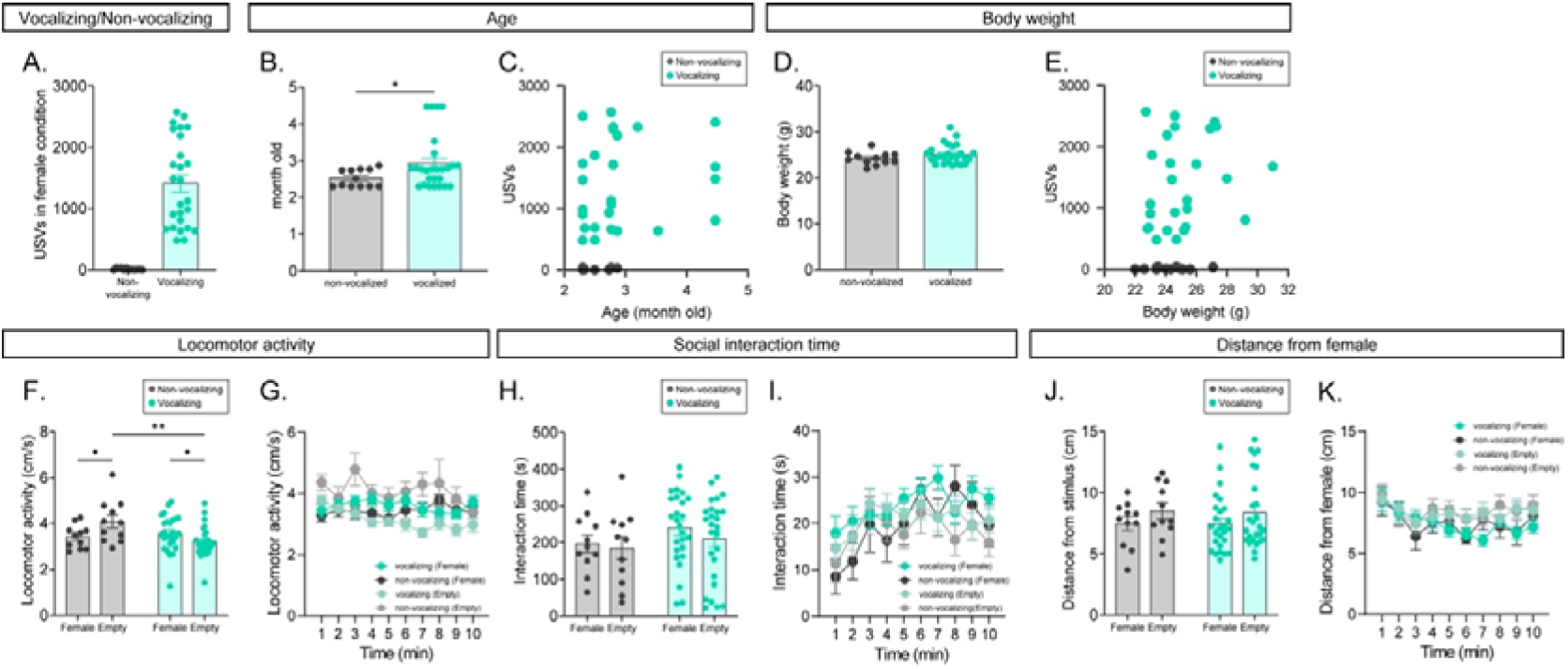
A. Number of USVs in the female condition in vocalizing and non-vocalizing mice. B. Age of vocalizing and non-vocalizing mice. C. Correlation between age and the number of USVs. D. Body weight of vocalizing and non-vocalizing mice. E. Correlation between body weight and the number of USVs. F. Locomotor activity in vocalizing and non-vocalizing mice. G. Time course of locomotor activity in vocalizing and non-vocalizing mice. H. Social interaction time in vocalizing and non-vocalizing mice. I. Time course of social interaction time in vocalizing and non-vocalizing mice. J. Distance from the female stimulus in vocalizing and non-vocalizing mice. K. Time course of distance from the female stimulus in vocalizing and non-vocalizing mice.

**Figure S2.**
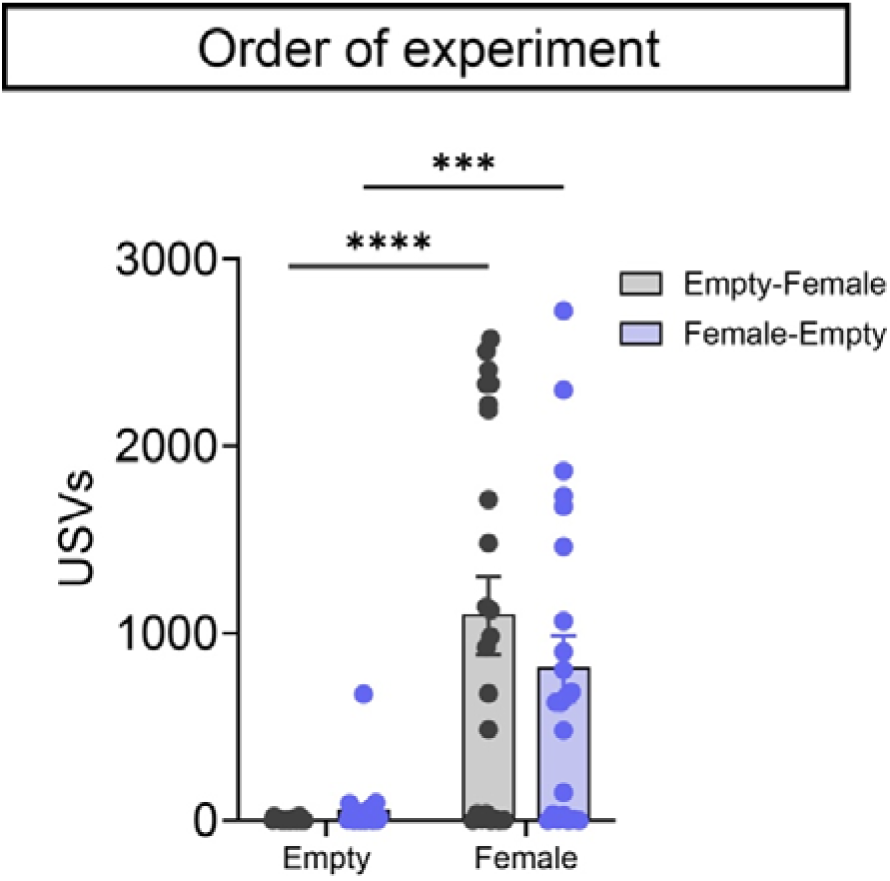
Number of USVs in the female and empty conditions, separated by stimulus presentation order (empty–female vs. female–empty).

### 3.2. Experiment 2

In Experiment 2, the vocalizing subjects from Experiment 1 were subsequently assigned to two groups and received either saline or lipopolysaccharide (LPS) 24 h prior to testing (Figure 4A). There were no significant differences in age or body weight prior to injection between the Saline and LPS groups (Figure S2). As expected, LPS administration significantly reduced body weight (Figure 4B; *t*(12) = 9.41, *p* < .0001, paired t test). Intraperitoneal injection of LPS also significantly reduced the number of USVs (Figure 4C; drug × stimulus interaction: *F*(1,24) = 13.62, *p* = 0.0011). Post hoc Fisher’s LSD tests revealed that LPS significantly decreased the number of USVs in the female stimulus condition (*p* < 0.0001). In addition, within the saline-treated control group, the number of USVs was significantly higher in the female stimulus condition than in the empty-basket condition (*p* < 0.0001). Time-resolved analysis further demonstrated that the number of USVs in the saline-treated group was consistently higher than that in the LPS-treated group throughout the testing session (Figure 4C). In contrast, LPS-treated mice exhibited markedly suppressed USV emission across all time bins, indicating a sustained reduction in vocal output over time (Figure 4D). Spatial analysis revealed that, whereas saline-treated mice emitted a high number of USVs when located in close proximity to the female stimulus, LPS-treated mice showed little to no USV emission even when positioned near the female stimulus (Figure 4E). These results indicate that LPS administration profoundly attenuated female-associated vocal communication, irrespective of temporal progression or spatial proximity to the stimulus. Furthermore, analysis of USV subtype composition revealed significant differences in the distribution of specific USV categories between the saline- and LPS-treated conditions. In particular, the proportion of up-frequency modulation (up-fm) calls was significantly reduced in the LPS-treated group compared with the saline-treated group (Figure 5F; main effect of classification: *F*(1.458, 22.54) = 17.03, *p* < 0.0001; drug × classification interaction: *F*(10,230) = 7.497, *p* < 0.0001; two-way ANOVA followed by Šídák multiple-comparison test). Consistent with the findings from Experiment 1, further analyses indicated that the distribution of USV categories was not modulated by either elapsed time during the session or spatial proximity to the female stimulus (Figure 4G-J). Overall locomotor activity during the social interaction task was significantly reduced following LPS treatment (Figure 4K; drug × stimulus interaction: *F*(1,24) = 11.35, *p* = 0.0025). Post hoc Fisher’s LSD tests revealed that LPS-induced reductions in locomotor activity were observed in both the female stimulus condition (*p* < 0.0001) and the empty-basket condition (*p* < 0.0001). In addition, within the saline-treated control group, locomotor activity was significantly higher in the empty-basket condition than in the female stimulus condition (*p* = 0.0022). The total duration of social interaction was also significantly decreased following LPS administration (Figure 4M; main effect of drug: *F*(1,24) = 19.49, *p* = 0.0002; two-way ANOVA), with significant reductions observed in both the female stimulus condition (*p* < 0.0001) and the empty-basket condition (*p* = 0.0010). The distance from stimulus was significantly decreased following LPS administration as well (Figure 4O, P; main effect of drug: *F*(1,24) = 24.4, *p* < 0.0001; two-way ANOVA), with significant reductions observed in both the female stimulus condition (*p* < 0.0001) and the empty-basket condition (*p* = 0.0064). Time-course analysis of behavior during the social interaction task further indicated that saline-treated control mice consistently were more active (Figure 4L), spent more time interacting with the female stimulus (Figure 4N), and remained closer to the stimulus (Figure 4P) throughout the session compared with LPS-treated mice.

**Figure 4.**
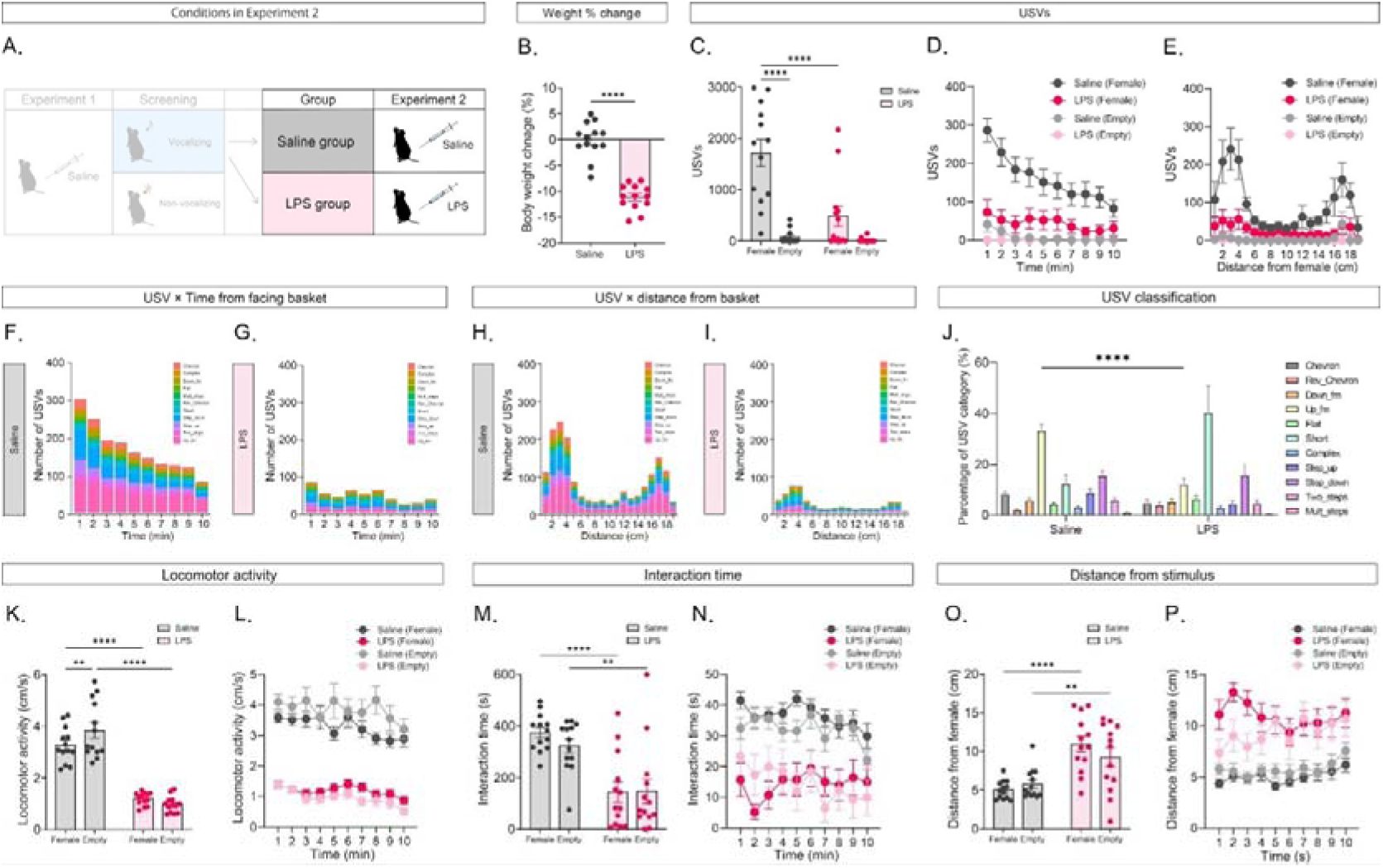
A. The vocalizing subjects were subsequently assigned to two groups, receiving either saline or lipopolysaccharide (LPS) in Experiment 2. B. Effects of LPS administration on the body weight after 24 hours. C. Effects of the LPS administration on the number of USVs under the female and the empty condition. D. Effects of LPS administration on the time course of the number of USVs. E. The number of USVs at the different distances from the basket. F. Distribution of USV categories. G. Time course of the distribution of USV categories under the female condition in the LPS group. H. Time course of the distribution of USV categories under the female condition in the saline group. I. Distributions of USV categories at different distances from the female in the LPS group. J. Comparison of the distribution of USV categories between the two groups. K. Comparison of locomotor activity between the two groups under the female and the empty condition L. Time course of the locomotor activity. M. Comparison of social interaction time between the two groups under the female and the empty condition. N. Time course of the social interaction time. O. Comparison of the distance from the basket between two groups under the female and the empty condition. P. Time course of the distance from the basket. Q. Error bars indicate the standard error of the mean, \**p* < 0.05, ** *p* < 0.01, **** *p* < 0.0001, N = 13.

**Figure 5.**
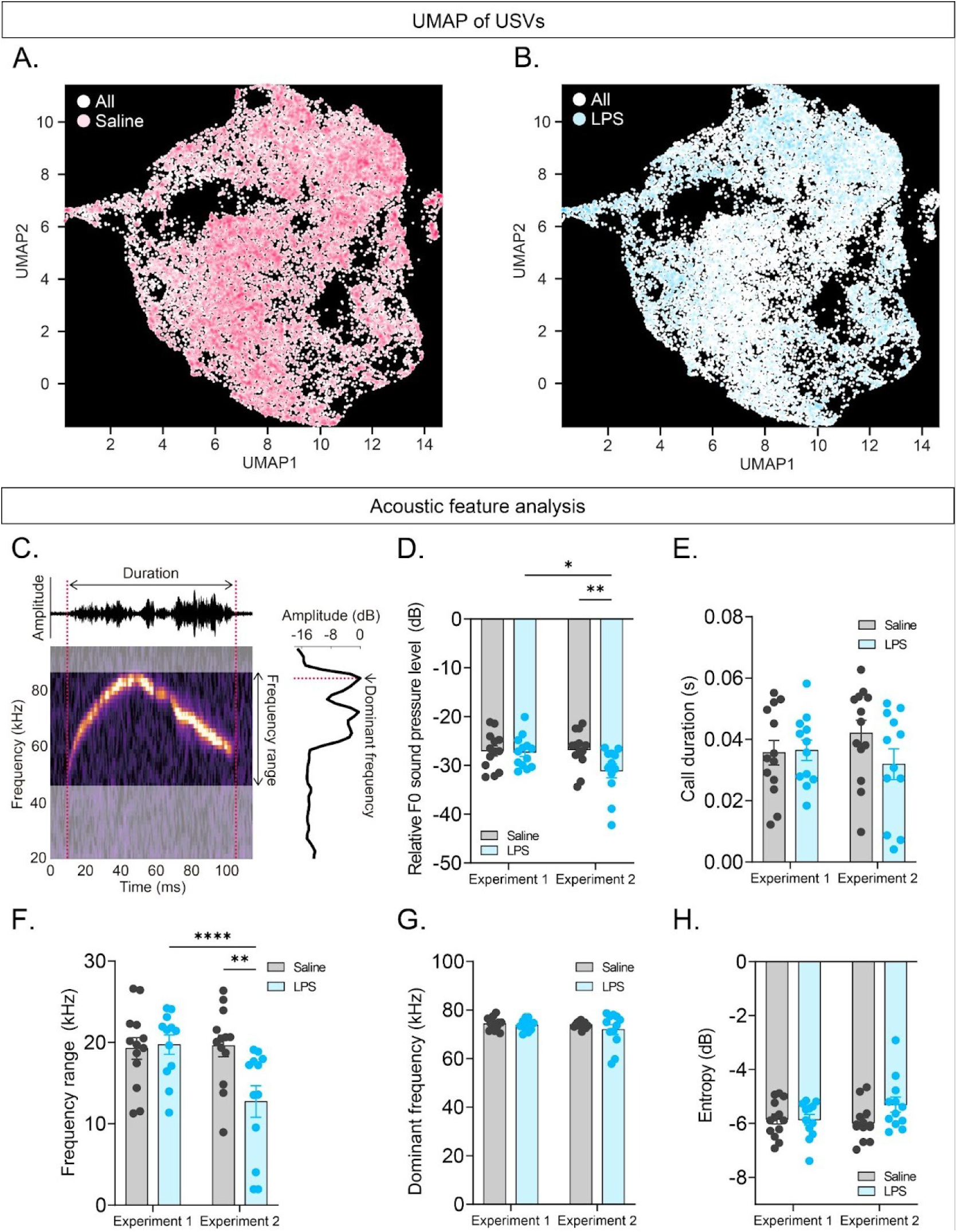
A. UMAP of saline condition. B. UMAP of LPS condition. C. Details of acoustic feature analysis. D. Effects of LPS on relative F0 sound pressure. E. Effects of LPS on call length. F. Effects of LPS on frequency range. G. Effects of LPS on dominant frequency. H. Effects of LPS on entropy. Error bars indicate the standard error of the mean. N = 13.

To characterize changes in USV acoustic structure, we applied dimensionality reduction using a variational autoencoder, an unsupervised learning approach (Goffinet et al., 2021). This analysis revealed a distinct shift in the distribution of acoustic features of USVs following LPS administration (Figure 5A,B), indicating that LPS altered the overall acoustic feature space of vocalizations. Furthermore, acoustic features of USVs, including sound pressure, call length, delta frequency, dominant frequency, and entropy (Figure 5B-H), in the female stimulus condition were analyzed using a mixed-effects model with Experiment (Experiment 1 vs. Experiment 2) as a within-subject factor and Treatment (saline vs. LPS) as a between-subject factor, with subject included as a random effect. LPS administration significantly reduced relative F0 sound pressure (Figure 5B; drug × time interaction: *F*(1,23) = 4.426, *p* = 0.0465). Post hoc Fisher’s LSD tests revealed that LPS significantly decreased relative F0 sound pressure after injection compared with pre-injection levels (*p* = 0.012) and relative to the saline-treated group (*p* = 0.0091). LPS administration also significantly reduced delta frequency (Figure 5F; drug × time interaction: *F*(1,23) = 12.56, *p* = 0.0017). Post hoc Fisher’s LSD tests confirmed that delta frequency was significantly decreased after LPS injection compared with pre-injection levels (*p* < 0.0001) and compared with the saline-treated group (*p* = 0.0018). In entropy, dominant frequency, or call length, there were no significant ef between the saline and LPS groups (entropy: *F*(1,23) = 1.773, dominant frequency: *F*(1,23) = 1.258, call length: *F*(1,23) = 0.845) or between the two experiments (entropy: *F*(1,23) = 1.348, dominant frequency: *F*(1,23) = 1.118, call length: *F*(1,23) = 0.091).

## 4. Discussions

In the present study, we demonstrated that LPS-induced systemic inflammation markedly attenuates male-female communication in mice. LPS administration reduced both the number of USVs and the duration of approach behavior toward females. In addition, the proportions of specific USV categories and several acoustic parameters, including sound pressure, were altered in LPS-treated mice compared with controls. These findings indicate that systemic inflammation affects not only the quantity but also the qualitative structure of courtship-related vocalizations. Given that USVs are widely regarded as a reliable indicator of social and reproductive motivation in rodents, the present results extend the concept of sickness behavior beyond its established effects on locomotion and general activity to include disruptions in social communication.

Importantly, the suppression of USV emission following LPS administration could not be fully explained by generalized reductions in locomotor activity or physical proximity to the female stimulus. Although LPS-treated mice exhibited decreased locomotion and reduced interaction time, the reduction in USVs persisted even when animals were located near the female and across all time bins. This dissociation suggests that inflammation-induced impairments in vocal communication are not merely secondary to reduced movement or engagement but instead reflect a more specific disruption of motivational or communicative processes underlying courtship behavior. Consistent with this interpretation, previous studies have reported that systemic inflammation impairs social motivation and interaction in rodents (Charoensaensuk et al., 2024; Fishkin & Winslow, 1997; Loh et al., 2023; Henry et al., 2008), yet most have relied on gross behavioral measures, such as proximity or locomotion (Hennessy et al., 2013; Savitz, 2019). By incorporating detailed analyses of USV output, the present study provides a more fine-grained assessment of social communication deficits induced by inflammatory states. In addition, our initial screening revealed that a subset of mice did not emit USVs despite showing comparable levels of approach behavior toward the female stimulus as those of vocalizing mice. Specifically, social approach behavior, body weight, and age did not differ between vocalizing and non-vocalizing mice. This finding suggests that USV emission is not strictly required for basic social interaction in this paradigm, possibly because mice were able to directly interact with the female through the wire mesh basket. Considering these points, USVs may serve as a specialized mode of social communication rather than a general indicator of social engagement. This interpretation further supports the idea that inflammation-induced reductions in USVs reflect a specific impairment in communicative processes rather than a general suppression of social behavior.

Beyond overall suppression, systemic inflammation selectively altered the composition and acoustic structure of male courtship USVs. In particular, reductions in up-frequency modulation calls and changes in sound pressure and delta frequency indicate that LPS affects not only the initiation of vocal output but also the fine control of vocal production. Previous work has suggested that distinct USV subtypes convey different motivational or affective states during male–female interactions (Chabout et al., 2015; Warren et al., 2018). From this perspective, inflammation-induced shifts in call composition and acoustic features may reflect altered internal states related to arousal, motivation, or affect, rather than a uniform suppression of vocal behavior.

Although the present study did not directly investigate underlying mechanisms, the observed effects of systemic inflammation on courtship USVs likely reflect the interaction of multiple neurobiological processes. Pro-inflammatory cytokines induced by LPS can influence brain function either directly, via circumventricular organs, or indirectly through vagal afferent signaling, thereby modulating activity in regions implicated in sociosexual communication, including the medial preoptic area, amygdala, and periaqueductal gray (Tschida et al., 2019; Michael et al., 2020; Xiao et al., 2023; Loh et al., 2023). In addition, inflammation-associated alterations in dopaminergic and serotonergic signaling have been reported in other behavioral contexts (Saggu et al., 2025; Lucido et al., 2021) and may contribute to reduced motivational drive for courtship behaviors. Furthermore, changes in respiratory or laryngeal motor control during inflammatory states could potentially underlie the observed modifications in acoustic features of USVs (Park et al., 2024). Taken together, these findings from prior studies provide a plausible interpretative framework, rather than direct mechanistic evidence, for understanding how systemic inflammation influences vocal communication.

An important consideration in interpreting the present findings is the timing and dose of LPS administration, as the behavioral consequences of systemic inflammation are highly dependent on these parameters. Acute LPS administration at doses comparable to those used here is well known to induce transient sickness behaviors, including reduced locomotion, social withdrawal, and motivational suppression (Kong et al., 2021; Morimoto et al., 2023; Yin et al., 2023). By contrast, relatively little is known about how systemic inflammation affects vocal communication. The present results demonstrate that even a single, acute inflammatory challenge is sufficient to disrupt courtship-related USVs, suggesting that vocal communication may represent a particularly sensitive and context-dependent target of systemic inflammation. Future studies directly comparing different doses, time points, and experimental models will be essential to determine whether these effects are specific to acute sickness states or generalize across broader inflammatory conditions.

Notably, the application of an unsupervised variational autoencoder revealed a global reorganization of the acoustic feature space following LPS administration. Unlike analyses focused on predefined acoustic parameters, this data-driven approach captures higher-order relationships among multiple vocal features, providing an integrative view of how inflammation reshapes vocal communication. The observed shift in latent acoustic representations suggests coordinated, system-level changes in vocal output, consistent with a systems-level disruption of vocal motor control and affective signaling. Such approaches may prove particularly useful for detecting subtle or distributed changes in communication patterns associated with disease states.

These findings have broader translational implications for understanding how systemic inflammation influences socio-communicative behaviors across species. In humans, inflammatory states are frequently associated with reduced social engagement and anhedonia, as observed in conditions such as major depressive disorder and chronic inflammatory diseases (Eisenberger et al., 2017; Yamamoto et al., 2026). Patients with depression also exhibit measurable changes in speech production, including reduced prosodic variation and altered vocal intensity, which have been proposed as objective biomarkers for diagnosis and treatment monitoring (Menne et al., 2024). In this context, inflammation-induced alterations in rodent USVs may provide a translational model for dissecting immune-related mechanisms underlying vocal and communicative changes in humans.

From a psychological perspective, these findings resonate with emerging views that social withdrawal and communicative blunting during sickness are not passive consequences of fatigue but actively regulated behavioral states. In humans, inflammatory challenges have been shown to reduce social motivation, alter emotional expression, and dampen vocal prosody (Eisenberger et al., 2017; Moieni et al., 2015). The present study suggests that similar immune-mediated mechanisms may shape the acoustic structure of social communication at a fundamental level. By linking peripheral immune activation to measurable changes in vocal signaling, this work provides a translational t-framework for investigating how inflammation contributes to social and communicative symptoms observed in psychiatric and neuroimmune disorders.

An important direction for future research will be to determine whether the observed changes in USVs are reversible and whether they can be selectively rescued by targeting inflammatory or neuromodulatory pathways. For example, pharmacological interventions that attenuate cytokine signaling or restore monoaminergic tone may help dissociate motivational deficits from motor impairments. In addition, incorporating neural recordings or circuit-specific manipulations during social vocalization tasks would provide critical insight into how immune signals interact with vocal communication networks in the brain.

A subset of mice did not emit USVs in Experiment 1 and were therefore excluded from subsequent USV-based analyses. Although all animals belonged to the same inbred strain and were reared under identical conditions, the factors underlying this marked individual variability in vocal output remain unclear. Importantly, vocalizing and non-vocalizing mice did not differ in body weight or in measures of social approach behavior toward the female stimulus, indicating that the absence of USVs does not necessarily reflect reduced social responsiveness. Vocalizing mice were older than non-vocalizing mice; however, the number of USVs was not significantly correlated with age, suggesting that age alone does not account for variability in vocal output. Moreover, this age difference may have arisen from uncontrolled factors such as cage-related effects or cohort differences and therefore should be interpreted with caution. Future studies will be required to clarify the potential contribution of age and related variables to individual differences in vocal communication. Such variability may reflect differences in motivational state, sensory processing, subtle prior social experiences, or unmeasured neurobiological traits, but these possibilities were not directly addressed here. Identifying the mechanisms that determine whether and when males engage in vocal communication remains an important open question.

Several limitations of the present study should be acknowledged. First, the use of a single LPS dose and a fixed post-injection interval limits our ability to determine the dose- and time-dependent effects of systemic inflammation on ultrasonic vocalizations. Future studies should examine a broader range of doses and time points to capture the full temporal dynamics of these changes. Second, while USVs provide a sensitive and ethologically relevant measure of social communication, they represent only one component of the complex behavioral repertoire underlying male-female interactions. Assessments of additional social and reproductive behaviors, as well as female responses, would provide a more comprehensive understanding of the impact of sickness states. Third, the present experiments did not directly investigate the neural or molecular mechanisms mediating LPS-induced changes in USVs and approach behaviors. Combining behavioral assays with neurophysiological, neurochemical, or neuroimaging approaches will be essential for identifying the specific brain circuits and signaling pathways involved. Finally, as this study was conducted exclusively in male mice, it remains unclear whether similar effects occur in females or in other social contexts, such as same-sex interactions. Addressing these limitations will be critical for translating these findings to broader models of inflammation-induced social dysfunction.

## 5. Conclusion

In summary, our study demonstrates that systemic inflammation induced by LPS profoundly disrupts male-female social interaction in mice, altering both the quantity and qualitative structure of courtship-related ultrasonic vocalizations. These effects likely reflect immune-mediated changes in neural circuits governing social motivation and communication, extending the concept of sickness behavior beyond its traditional boundaries. By establishing a robust link between inflammation and altered socio-communicative behaviors, our findings provide a foundation for future mechanistic and translational research aimed at mitigating the social deficits associated with inflammatory states in both preclinical models and human conditions.

## CRediT authorship contribution statement

**Mizuki Yamamoto**: Writing – review & editing, Writing – original draft, Validation, Methodology, Investigation, Formal analysis, Conceptualization. **Haruki Kasahara**: Validation, Methodology, Investigation, Formal analysis. **Kazuko Hayashi**: Resources, Validation, Funding acquisition. **Yuta Tamai**: Writing – review & editing, Writing – original draft, Resources, Validation, Methodology, Formal analysis, Funding acquisition. **Koji Toda**: Writing – review & editing, Writing – original draft, Validation, Resources, Methodology, Investigation, Formal analysis, Supervision, Resources, Funding acquisition, Conceptualization

## Ethical statement

The experimental and housing protocols adhered to the Japanese National Regulations for Animal Welfare and were approved by the Animal Care and Use Committee of the Keio University.

## Acknowledgement

We thank Hiroto Inoue, Shuntaro Matsushima, Yuuki Fujii, and Rei Takahashi for their assistance on animal care.

## Abbreviations

F0: initial frequency of fundamental frequency
LPS: lipopolysaccharide
TLR4: Toll-like receptor 4
UMAP: uniform manifold approximation and projection
USVs: ultrasonic vocalizations

## Data availability

Data supporting the findings of this study are available from the corresponding author upon reasonable request. The original codes written for the analysis are available from the corresponding author upon reasonable request.

