## Supplementary information for "Lipopolysaccharide-induced systemic inflammation alters courtship ultrasonic vocalizations in male mice"

Supplementary material

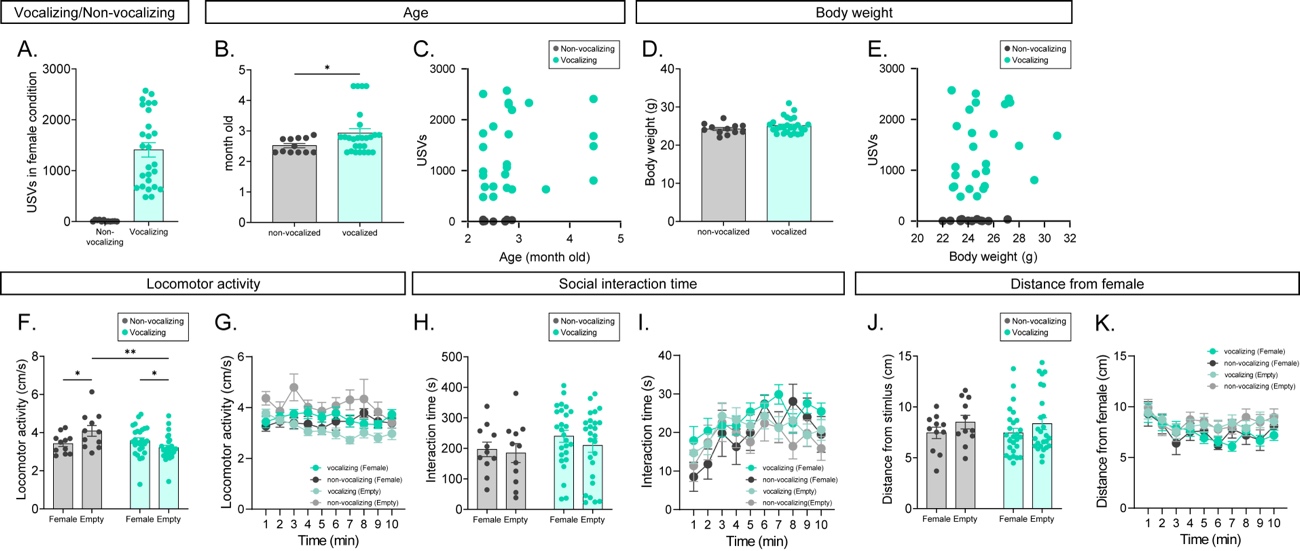

Figure S1.

A. Number of USVs in the female condition in vocalizing and non-vocalizing mice. B. Age of vocalizing and non-vocalizing mice. C. Correlation between age and the number of USVs. D. Body weight of vocalizing and non-vocalizing mice. E. Correlation between body weight and the number of USVs. F. Locomotor activity in vocalizing and non-vocalizing mice. G. Time course of locomotor activity in vocalizing and non-vocalizing mice. H. Social interaction time in vocalizing and non-vocalizing mice. I. Time course of social interaction time in vocalizing and non-vocalizing mice. J. Distance from the female stimulus in vocalizing and non-vocalizing mice. K. Time course of distance from the female stimulus in vocalizing and non-vocalizing mice.

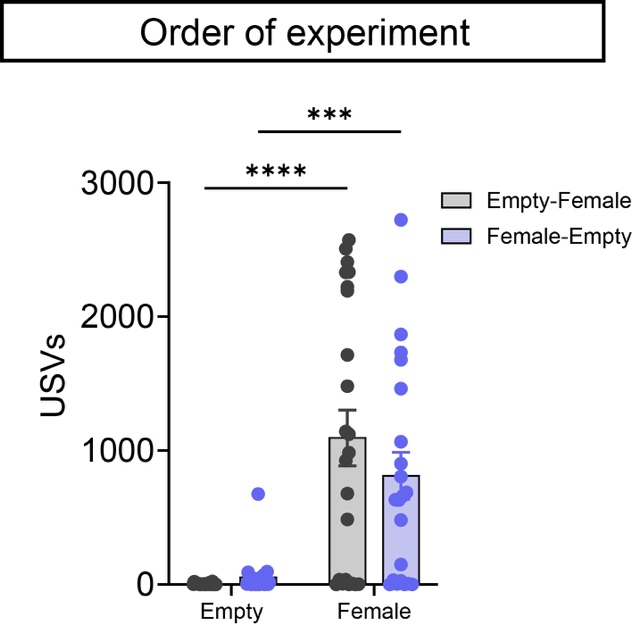

Figure S2.

Number of USVs in the female and empty conditions, separated by stimulus presentation order (empty–female vs. female–empty).
